# Immunosuppressive regimens based on Cyclophospamide or Calcineurin inhibitors: Comparison of their effect in the long term outcome of Primary Membranous Nephropathy

**DOI:** 10.1101/630525

**Authors:** Maria Stangou, Smaragdi Marinaki, Evangelos Papachristou, Chrysanthi Kolovou, Erasmia Sambani, Synodi Zerbala, Panagiota Papadea, Olga Balafa, Karolos-Pavlos Rapsomanikis, Aimilios Andrikos, Panagiota Manolakaki, Dorothea Papadopoulou, Efstathios Mitsopoulos, Paraskevi-Eyh Andronikidi, Vasiliki Choulitoudi, George Moustakas, Dimitra Galitsiou, Eugene Dafnis, Constantinos Stylianou, Ioannis Stefanidis, Spyridon Golfinopoulos, Stylianos Panagoutsos, Maria Tsilivigkou, Apostolos Papadogianakis, Ioannis Tzanakis, Athanasios Sioulis, Dimitrios Vlachakos, Eirini Grapsa, Sophia Spaia, Nikolaos Kaperonis, Christos Paliouras, Christos Dioudis, Fani Papoulidou, Theofanis Apostolou, Christos Iatrou, Ioannis Boletis, Dimitrios Goumenos, Aikaperini Papagianni

## Abstract

Management of the Primary Membranous Nephropathy (PMN) usually involves administration of immunosuppressives. Cyclophosphamide (Cyclo) and Calcineurin Inhibitors (CNIs) are both widely used but only limited data exist to compare their efficacy in long term follow-up.

**Aim:** of the present study was to estimate and compare long term effects of Cyclo and CNIs in patients with PMN.

**Patients-Methods:** Clinical data, histologic findings and long term outcome were retrospectively studied. The response to treatment and rate of relapse was compared between patients treated with CNIs or Cyclo based immunosuppressive regimens.

**Results:** Twenty three centers participated in the study, with 752 PMN patients (Mean age 53.4(14-87)yrs, M/F 467/285), followed for 10.1±5.7 years. All patients were initially treated with Renin Angiotensin Aldosterone System inhibitors (RAASi) for at least 6 months. Based on their response and tolerance to initial treatment, patients were divided into 3 groups, group I with spontaneous remission, who had no further treatment, group II, continued on RAASi only, and group III on RAASi+immunosuppression. Immunosuppressive regimes were mainly based on CNIs or Cyclo. Frequent relapses and failure to treatment were more common between patients who had started on CNIs (n=381) compared to those initially treated with Cyclo (n=110), relapse rate: 25.2% vs. 6.4%, p<0.0001, and no response rate: 22.5% vs. 13.6%, p=0.04, respectively.

**Conclusions:** Long term follow up showed that administration of Cyclo in PMN is followed by better preservation of renal function, increased response rate and less frequent relapses, compared to CNIs.

## Introduction

Membranous Nephropathy (MN) is the most common cause of nephrotic syndrome in adults, with an overall global incidence reaching to 1.2 per 100,000 per year, as estimated in a recent review covering the period 1980-2010 [1,2]. Outcome of disease is variable, ranging from spontaneous remission to progressive renal failure reaching end stage renal disease [1-4]. Undoubtedly, after the discovery of innovative pathogenic antibodies, the M-type phospholipase A2 receptor (PLA2R), we are on the way to changing our aspect on the disease, in terms of research and clinical management [5-7]. However, critical questions regarding diagnostic approach and follow up, assessment of renal biopsy findings, and most importantly, selecting and applying the best therapeutic regime for each patient still remain.

Treatment of the disease has been a challenge for many years. KDIGO guidelines was a remarkable attempt to set treatment rules. Therefore, they recommend the use of Cyclo, in the six month cycle Ponticelli regime, or alternatively, CNIs as first line immunosuppressive treatment in patients who had not responded to RAASi for at least six months, or in patients with rapidly declining renal function or those with life threatening complications due to proteinuria [4]. Both treatment options have their own side effects and both have undergone alterations and improvements in order to minimize complications [1,4].

Although Cyclo and CNIs have been used in the treatment of PMN for more than twenty years, there are not enough data to support one against the other, and the few prospective studies have only a short term follow up, up to 12 months, which is not a long enough period to come to confident conclusion [8-12]. Aim of the present retrospective multicenter study, was to evaluate the long term effects of Cyclo and CNI treatments in a large cohort of PMN patients.

## Patients-Methods

**Inclusion criteria were:** 1. Biopsy proven MN, 2. Exclusion of secondary causes, 3. Initial treatment with RAASi continued for at least 6 months, 4. Follow up for at least 12 months, unless patients reached end stage renal disease (ESRD) or died

## Presentation, treatment and follow up

Twenty three centers collaborated in this study. Diagnosis of MN was based on renal biopsy findings, and all information, including clinical symptoms, medication, past medical history, histology, laboratory results at time of diagnosis, routine investigation to exclude secondary causes, response to treatment, relapses and outcome were collected from the Greek Registry of Membranous Nephropathy, part of Glomerular Diseases Network, under the auspices of the Hellenic Society of Nephrology.

e-GFR was estimated based on MDRD equation. Definitions of remission (complete, partial, no remission) were adopted from KDIGO. Relapse of the disease was defined as the re-appearance of nephrotic syndrome after achieving complete or partial remission.

Outcome of the disease at the end of follow up was estimated by the primary end point, defined as no-response or ESRD. The presence of two or more relapses during follow up was defined as secondary end point.

End of follow up was considered the last visit to outpatients clinic or for those who reached ESRD or died, the time when they started on dialysis method or day of death.

## Treatment protocols

According to patient records, the initial and subsequent treatment protocols were reported, in addition to response to therapy, defined as total, partial or no remission, number of relapses, need to change treatment protocol, different treatment protocols and final outcome. Treatment decisions were made by treating physicians and were based to KDIGO guidelines, histology, co-morbidities and the experience of each center. All patients started on treatment with RAASi. After at least 6 months trial with RAASi, immunosuppressive treatment was added to those, who had partial or no response. A small group of patients were started on immunosuppression earlier than 6 months, either because of rapidly deteriorated renal function and/or life threating complications due to the nephrotic syndrome. Finally, some patients were not offered treatment with immunosuppressives, either because of severely impaired renal function and/or histology, or because of increased probability of infections.

## Histology

A global review and evaluation of renal biopsy reports was performed, and the characteristics estimated included percentage of obsolescent glomeruli (defined as global sclerosis), presence or absence of focal segmental sclerosis, vascular hyalinosis, severity of tubular atrophy and interstitial fibrosis. The severity of tubular atrophy was rated in a scale of 0-2 based on the percentage of affected tubules (<25%, 25-50%, >50%, respectively); in a similar way the extension of interstitial fibrosis was semiquantively estimated and rated as 0, 1, 2 for absent-mild, moderate, severe interstitial fibrosis, respectively.

## Statistical analysis

Statistical analysis was performed using SPSS 23.0 software for Windows. P values of <0.05 were considered as statistically significant. Data from normally distributed variables were expressed as mean ± SD, and Student’s t test or ANOVA test was performed to compare differences between groups. Non–normally variables were expressed as medians and interquartile range (IQR), and differences between groups were compared by Mann–Whitney U or Kruskal-Wallis H test. Multivariate analysis was performed to estimate the independent parameters correlated with the outcome of renal function. Renal survival differences based on different treatment options were estimated with Kaplan-Meier test.

## Results

### Patients characteristics at time of diagnosis

The records of 1098 patients with MN, diagnosed during the period 1995-2015, were retrospectively studied. The whole cohort of patients went under thorough investigation, initially, to exclude secondary forms of MN, such as systemic diseases, infections, hematological diseases, and subsequently, to exclude patients who did not fill the inclusion criteria. After extensive analysis, 752 patients with PMN, who fulfilled the described criteria were included in the study (Figure 1).

**Figure 1.**
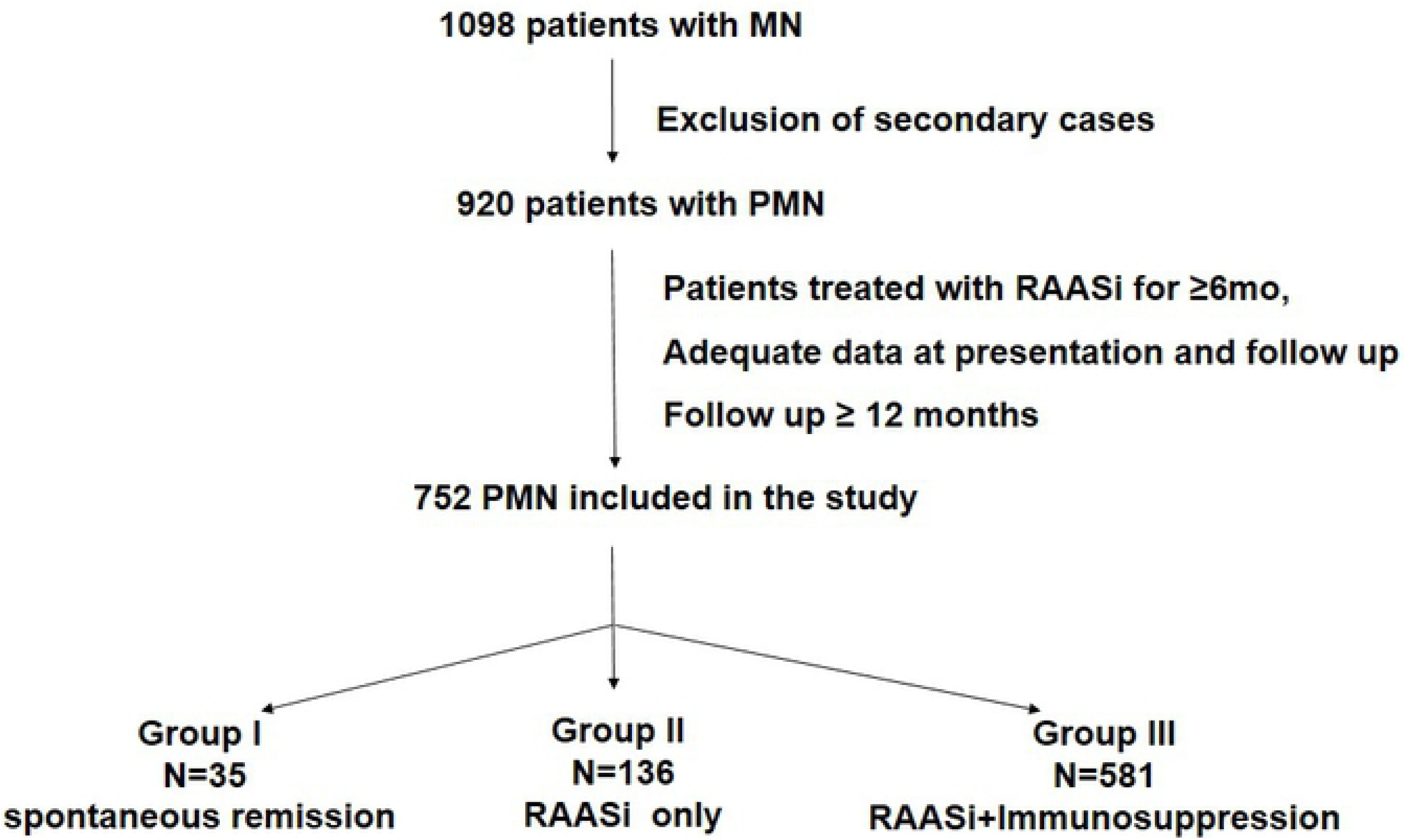

Clinical and laboratory data at the beginning and at the end of the study are shown on table 1. All patients were European Caucasians (94% Greek and 6% East European origin), with a mean age of 53.4±14years (15-85years) and most of them, 468/752 (62.2%), were men. At presentation, most patients were at stage II or III of chronic kidney disease (355/752, 47.2% and 203/752, 26.9% respectively). The vast majority presented with nephrotic syndrome (613/752, 81.5%), hypertension (466/752, 61.9%), hypoalbuminaemia, defined as serum albumin levels<3gr/dL, (481/752, 63.9%) and dyslipidaemia, defined as serum cholesterol >250mg/dl and/or serum triglycerides >160mg/dl, (482/752, 64%). Microscopic hematuria was present in 344/752 (45.7%).

Following their initial response to 6 months treatment with RAASi, patients were divided in 3 groups. **Group I (n-35)**: Patients who had a rapid response, probably spontaneous remission, and stopped RAASi after the initial 6 months, **Group II (n=136):** Patients who continued with RAASi as monotherapy, either because of remission or because they were not considered suitable for immunosuppression, and **Group III (n=581):** Patients with partial or no response to RAASi, who were subsequently treated with a combination of RAASi+Immunosuppression. Table 2 describes the clinical characteristics and histology of patients allocated to each treatment option, and also, their disease outcome. Follow up of the whole cohort of patients was 121.8 (14-372)months.

### Treatment and Outcome

#### A. Renin-Angiotensin-Aldosterone System inhibitors

RAAS inhibitors were initially given to all 752 patients for at least 6 months.

##### i) Spontaneous remission

Thirty five patients, 35/752 (4.65%), (Group I) had an early spontaneous remission; these patients were the youngest, and had mild histological lesions, regarding both glomeruli and tubulointerstitial compartment. After a follow up of 128.5 (34-336) months, eGFR and Uprot were reduced but not significantly (Z=-1.1, p=NS and Z=-1.8, p=NS respectively). In this group of patients outcome of renal function, was correlated only with eGFR at time of diagnosis and the degree of interstitial fibrosis (Table 3).

##### ii) Renin-Angiotensin-Aldosterone System inhibitors as monotherapy

One hundred thirty six patients, 136/752 (18.09%), (Group II) continued on RAASi only. These patients were the oldest, with severe renal function impairment and chronic histological lesions. Their follow up was 101.4 (14-312) months; during this period, eGFR and Uprot were reduced significantly (Z=-3.2, p=0.001 and Z=-6.6, p<0.0001 respectively). Final eGFR levels correlated with age, renal function at presentation and severity of tubuloinerterstitial lesions (Table 2, 3).

#### B. Immunosuppressive treatment

Patients in this group III, n=581 (77.26%), had the higher levels of proteinuria at presentation and less severe histology compared with group II. After 115.5 (14-372) months follow up, the levels of eGFR had positive correlation with age, renal function and proteinuria at presentation, and also, with degree of global sclerosis, presence of FSGS and vascular hyalinosis and severity of tubulointerstitial lesions (Table 3). Multivariate analysis revealed that independent factors correlated with response to treatment were the degree of initial proteinuria (p<0.0001), percentage of obsolescent glomeruli (p<0.0001) and severity of interstitial fibrosis (p=0.001).

In 491/752 Immunosuppressive regimens used were either based on CNIs (as monotherapy, or in combination with steroids) or on cyclophosphamide (per os or in modified Ponticelli regime). Seventy four patients received other immunosuppressive treatments, including Ponticelli regime (with chlorambucil), Mycofenolate Mofetil, Azathioprine or Rituximab, however, the effect of these regimens was not analyzed, as in the present study analysis was performed for patients who received CNI-based or Cyclo-based immunosuppressive treatment schemes.

##### i) CNIs or Cyclo used as initial treatment schemes

###### CNIstart Group

Initial immunosuppressive therapy was based on CNIs (CNIstart Group) in 381/581 (65.6%) and on Cyclo (Cyclostart Group) in 110/581 patients (18.9%).

Patients in CNIstart Group (n=381, M/F: 244/137) were 53.8±15 years old, and, after a follow up period of 114 (15-336)months, their renal function changed as follows: Screat increased from 1.01±0.5 to 1.6±1.5mg/dl, p<0.0001, eGFR reduced from 70.2±22 to 55.1±24ml/min/1.73m^2^, p<0.0001 and Uprot from 7.9±5.2 to 2.2±0.2gr/24hr, p<0.0001.

Histology showed FSGS in 258/381 (67.7%) tubular atrophy graded as 0, 1 and 2 in 181/381 (47.5%), 180/381 (47.3) and 20/381 (5.2%), respectively, interstitial fibrosis 0, 1, 2 in 185/381 (48.6%), 170/381 (44.6%), and 26/381 (6.8%) respectively, and vascular hyalinosis in 185/381 (48.6%).

###### Cyclostart group

Patients on Cyclostart group (n=110, M/F: 78/32) were 52.9±13 years old, and, after 115 (15-372) months follow up, their renal function changed: Screat from 1.1±0.3 to 1.3±1.1mg/dl, p<0.0001, eGFR from 70.6±20.5 to 62.2±22ml/min/1.73m^2^, p<0.0001 and Uprot from 8.2±5.3 to 2.3±3gr/24hr, p<0.0001.

Histology showed presence of FSGS in 65/110 patients, (59%), tubular atrophy graded as 0 in 39/110 (35.5%), 1 in 66/110 (60%), 2 in 5/110 (4.5%), interstitial fibrosis 0 in 46/110 (41.8%), 1 in 58/110 (52.7%), 2 in 6/110 (5.5%) and vascular hyalinosis in 49/110 (44.5%).

CNIstart and Cyclostart group patients, had no significant differences in terms of renal function and histology at presentation.

Patients started on Cyclo had significantly better outcome (Figure 2A), and this was the same for the whole cohort of patients and for those presented with nephrotic syndrome (Figure 2C). Eighty six patients in CNIstart group (22.5%) and 15 (13.6%) in Cyclostart group reached the primary end point, p=0.04. Similarly, initial treatment with CNIs was followed by significantly more relapses, 96 (25.2%) patients in CNIstart group reached secondary end point, compared to 7 (6.4%) patients in Cyclostart group, p<0.0001 (Table 4). However, 31/110 (28.1%) patients who received Cyclo transferred to alternative treatment regimes during follow up, the reasons being no response, relapses and reluctance of treating physicians to administer more than 1 or 2 courses of Cyclo. This rate of changing treatment was significantly higher compared to CNIstart group (69/381, 18.4%) (Chi-square test 5.3, p=0.02).

**Figure 2.**
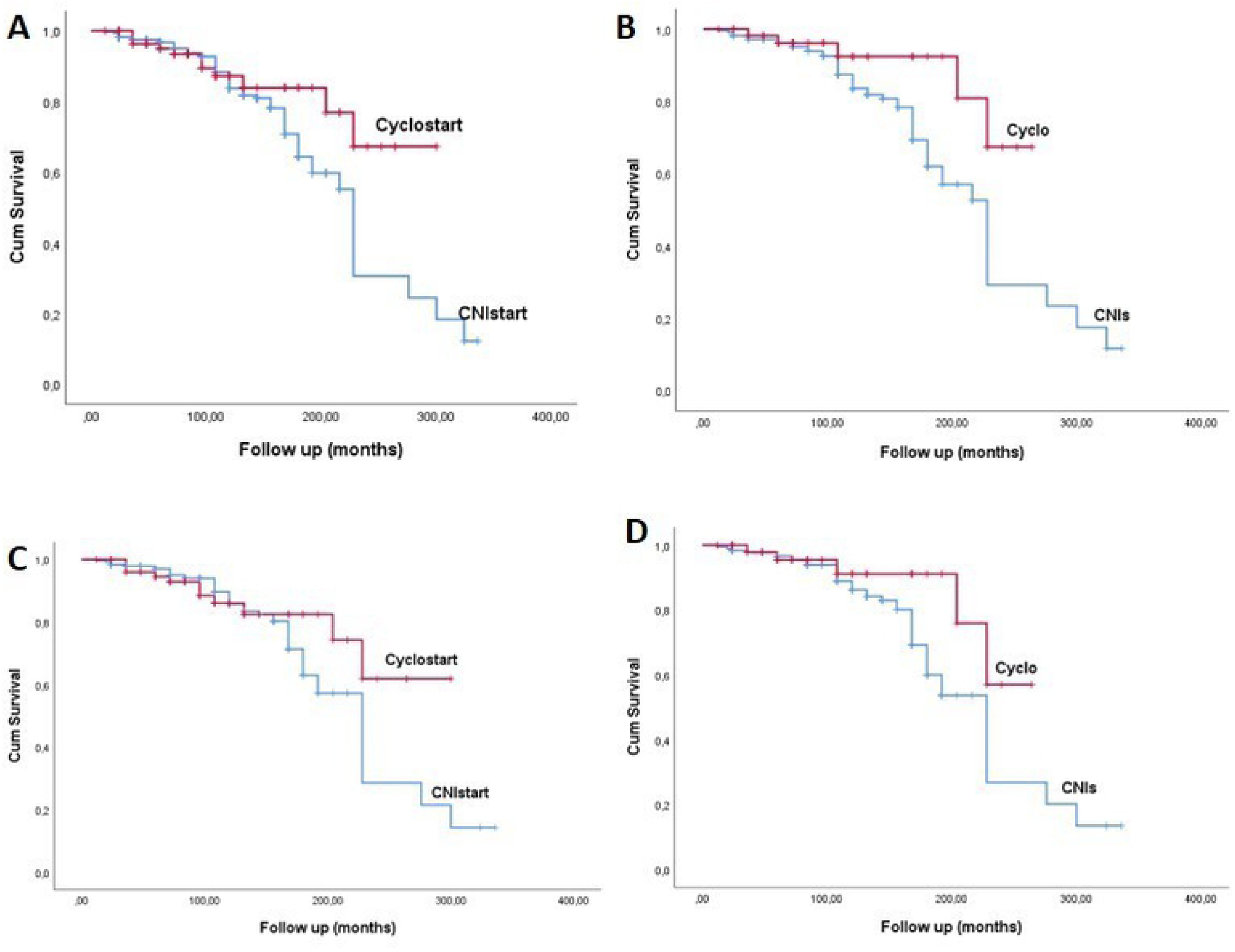

##### ii) CNIs or Cyclo based schemes as the only treatment

Three hundred and twelve patients from the CNIstart group (Age 54.5±15yrs, M/F 197/115) and 79 from Cyclostart group (Age 53.1±13.2yrs, M/F 55/24) did not receive any other immunosuppressive treatment during their follow up. Outcome of these patients are depicted on tables 5 and 6. Severity of proteinuria was reduced in both groups, as was renal function, which however at the end of the study was significantly worse in patients receiving CNI based treatment, 55.1±24.6 vs. 67.8±17.8ml.min/1.73m^2^, p=0.01. Patients treated with CNIs only, had significantly higher rates of no response and more frequent relapses, 76/312 vs. 6/79 reached primary end point respectively, p=0.0001, and 60/312 vs. 4/79 reached secondary end point respectively, p=0.002 (Table 6, Figure 2B, 2D).

### Independent parameters predicting outcome

In the whole cohort of patients, independent parameters predicting renal function outcome, were eGFR (R^2^=0.297, p<0.0001), levels of Uprot at presentation (R^2^=0.542, p<0.0001), the severity of tubular atrophy (R^2^=0.423, p<0.0001) and presence of FSGS (R^2^=0.507, p<0.0001), when eGFR at last follow up was used as the dependent variable. Similarly, when the response rate was used as dependent parameter, independent variables were degree of proteinuria (R^2^=0.16, p<0.0001), severity of tubular atrophy (R^2^=0.1, p=0.001) and presence of FSGS (R^2^=0.2, p<0.0001).

## Discussion

To our knowledge, this study, based on a database of 752 PMN patients, has included the largest number of patients with the longest follow up, reaching more than 10years.

Treatment protocols applied were based on KDIGO guidelines. We evaluated clinical data and histology of patients allocated to each of the three treatment alternatives after the 6 months course of RAASi, no treatment (Group I), RAASi only (Group II) and RAASi+Immunosuppression (Group III), and we tried to define the profile of patients who will benefit from each treatment. Furthermore, we estimated and compared the effect of CNIs or Cyclo, in a long term follow up period.

Clinical presentation in the vast majority of patients included nephrotic syndrome, hypertension mild to moderate renal function impairment and microscopic hematuria. Defining parameters which can correlate to renal function and guide treatment has always been an important issue in the clinical management of PMN [13-14]. Conflicting results about the importance of demographic and clinical data in renal function outcome, have led investigators to suggest a model of prediction and describe the Toronto Risk Score. The score, which was defined many years ago and simplified later, has an accuracy of 85-90% in identifying patients at risk of progression [15,16]. In our study, apart from the clinical presentation, we also evaluated renal biopsy findings. In agreement with previous studies, we found that severity of tubulointerstitial lesions play a central role in renal function outcome, but we also described that the percentage of obsolescent glomeruli, the presence of FSGS and vascular hyalinosis are also important in patients who need immunosuppressive treatment. To our knowledge, our study is the first one to emphasize the role of global sclerosis and FSGS in disease outcome, mainly in patients who receive immunosuppression.

Patients in whom proteinuria was improved after 6 month treatment with RAASi were either young, with mild histological lesions and preserved renal function, or old patients with chronic clinical and histological profile. Treatment with RAASi has been established in chronic proteinuric diseases. However, RAASi is not expected to have any effect in immune active disease, therefore, patients who will benefit are either at early non-nephrotic stage, or have progressed to advanced disease [17-21]. It is not easy to assess immunological status in PMN, although the recent finding of PLA2R-Ab can provide some information, as low levels usually represent inactive phase, predict beneficial outcome and thus, suggest treatment with RAASi and no immunosuppression [22].

Based on KDIGO and some recent studies, the percentage of patients who will finally start immunosuppressive treatment will be substantially reduced if immunosuppressive treatment is postponed for at least 6 months, or until mild deterioration of renal function. There has been a debate about this option, as patients who start immunosuppression at an early stage are more likely to have a rapid remission of nephrotic syndrome, although not always accompanied by improvement of renal function [4,23,24].

A common dilemma clinicians have to face is to choose between cyclophosphamide and CNIs. In most countries, Rituximab is used only when both these treatments have failed, besides, the optimal dose, short and long-time side effects are under investigation and no definite conclusions have been made so far. KDIGO guidelines suggest the use of either Cyclo or CNIs, but do not support one choice against the other, there are no exact indications, and both agents carry their own side effects. [25,26]. Cyclophospamide has effectively substituted for melphalane in “Ponticceli regime”, there have been several studies that support its use against melphalan and, also customized in the 6 month cycle Ponticelli protocol instead of per os continuing regimens [26,27]. CNIs have been used either in combination with steroids or as monotherapy, the main concerns being the risk of nephrotoxicity and increased rate of relapse after discontinuation of treatment [4, 28-30].

Few studies have compared the effect of alkylating agents with that of CNIs, in the treatment of PMN and initially showed a comparable efficacy in proteinuria reduction and rate of relapse [8-11]. However, all had a short term period, not exceeding 12 months, and they did not refer to outcome after discontinuation of treatment. There is only one randomized prospective study with extended follow up to 24 months, which confirmed previous results of similar and comparable effect of the two therapeutic modalities in the short term, but showed inferiority of Tacrolimus after 18 and 24 months of follow up [12,31]. Our study is the first one to compare these two treatment options in long follow up. Patients were matched for age, renal function, severity of proteinuria and histology. Our results revealed significantly higher rates of response, better preserved renal function and lower frequency of relapses in Cyclo treated patients, compared to CNIs. Even more importantly, these differences were consistent between patients who started with Cyclo or CNIs and subsequently transferred to other immunosuppressives; suggesting that the beneficial effect of Cyclo remains even after subsequent change of treatment. Surprisingly however, we fount that, although patients on CNIs were more likely to relapse or not respond, they were less likely to change therapeutic protocol. This is due to the fact that patients who have a relapse after CNIs reduction or withdrawal, do not change treatment but they increase the dose or restart the drug; consequently, a great proportion ends up continuing treatment with small doses of CNIs, which can probably avoid relapses, but may be implicated to more severe renal impairment, due to CNI nephrotoxicity. In contrast, patients who have frequent relapses or do not respond to treatment with Cyclo will have to change their initial protocol to alternative immunosuppressive regimens, as physicians are cautious with the cumulative dose of cyclophosphamide.

There are some limitations in our study; mainly regarding the reports of side effects. Although adverse events from immunosuppression were reported in the patients files, the exact number and rate of adverse events cannot be strictly estimated, as mild side effects were treated in local hospitals and not stated on the files.

However, this is the first study to compare the effect of the two most common treatment options in PMN, Cyclo and CNIs, in a large number of patients followed for more than 10 years. The use of cyclophosphamide based regimes as first choice of immunosuppressive treatment seems more promising for the long term outcome even if alternative immunosuppressive regimes will be subsequently used during follow up.

